# Correlates of Sleep and Arousal via Matrix Methods

**DOI:** 10.1101/2021.01.20.427445

**Authors:** Siamak K. Sorooshyari, Luis de Lecea

## Abstract

Conventional computational modeling of sleep and arousal are primarily brain-centric in restricting attention to data from the nervous system. While such a view is warranted, the importance of considering the coupling of peripheral systems in the causes and effects of sleep are being increasingly recognized. An analysis is presented that has the capability to incorporate neural recordings of different modalities as well as data from the metabolic and immune systems. We formulate a matrix-based approach for inference of the arousal state based on the activity level of cell types that will comprise the matrix components. While the presented computations are intended to predict sleep/arousal, it is anticipated that a scrutiny of the structure of the matrices will provide insight into the dynamics of the constituent systems. A model is also proposed to consider the interaction of the signals recorded across the neural, metabolic, and immune systems in leading to the arousal state.

## 1. Introduction

Sleep disturbances are a core symptom of most neuropsychiatric disorders; however, the neuronal underpinnings of such association are unknown. Despite the profoundly negative effects of such disorders on public health, progress in understanding the pathophysiology and the discovery of new therapeutic targets have been slow. A modeling of the brain’s complex, multidimensional responses and how multiple neural circuits regulate behavioral state transitions will benefit our understanding of brain function and disorders. However, defining and correlating brain state transitions with specific behavioral changes is challenging. For example, the transition between sleep and arousal is associated with brain-wide changes in intrinsic cellular, metabolic, and neuronal activity characteristics, which are linked with specific physiological and behavioral outputs. Attempts to understand the implications of the correlations are typically limited by experimental capabilities and the complex nature of the brain, thus, analytical techniques can set a hypothesis-driven framework.

Sleep is evolutionarily conserved across the animal kingdom and is regulated by circadian and homeostatic interactions [1]. The dominant paradigm for assessing sleep in mammals is measurement of cerebral cortex electrical activity using an electroencephalogram (EEG). EEG Signals combined with electromyogram (EMG) are often used to distinguish wake, rapid eye movement (REM), and non-REM (NREM) sleep states. Although EEG is a well-established marker of sleep in mammals [2][3], the specific role of the voltage fluctuations is not well understood as the activity generated by a sub-population of neurons cannot be differentiated from the context of entire brain region. Recent work on the demonstration of local sleep phenomena [4] and complex dynamics driven by homeostatic mechanisms [5] highlight the limitations of using EEG as the only measure to score vigilance. Monitoring brain states during sleep and wakefulness in mammals is also challenging because the mammalian brain is hidden within the opaque cranium and imaging activity at single cell resolution is restricted to small superficial regions. These limitations can be partially overcome by fiber photometry, a technique that allows recording calcium signals of cell-type specific populations in deep brain structures in behaving animals [1]. This method collects fluorescence from bulk activity generated by ensembles of neurons transduced with a genetically encoded calcium indicator. Although the central orchestrator, the brain is not the only entity determining or affecting arousal. Energy metabolism is naturally involved in arousal with the conservation and allocation of energy postulated as one of the main functions of sleep [6][7][8]. The metabolic and immune systems are both inducers and responders to the arousal state and the properties of sleep [9][10]. Increasingly, both systems are being considered along with neuronal recordings to monitor, understand, and predict arousal and the pathologies associated with sleep.

Brain recordings attained via EEG are frequently reduced to a spectrogram that provides time-frequency information that can be represented in matrix form. While EEG is effective in its portrayal of global cortical activity, the arousal system is quite heterogeneous. Thus, EEG recordings provide a partial and somewhat incomplete picture since the local network activity of sub-cortical regions cannot be fully captured. Throughout the circadian cycle the various brain regions and neural populations can exhibit activity or characteristics reflective of sleep/arousal while the animal’s (aggregate) brain state is sleep/wake. Recent developments in genetically encoded sensors have enabled recording calcium activity deeper and more precisely in a brain, thus allowing for the identification of brain states in various neural circuits while an animal passes through sleep/wake cycles [11]. A relatively recent advancement in neuroscience has been the aggregation of such concomitant, local activity measurements to gain predictive capability of brain state in subsequent epochs [12]. Analogous to EEG data, calcium recordings can be represented in matrix form when considering multichannel activity recorded from different populations across a recording interval.

Intrinsically linked to arousal, immune system and metabolic data can also be represented in matrix form in order to extract structure on their contribution to a brain state. In particular, metabolic data is expected to yield fewer time points than the neuronal recordings mentioned, however, aptamers and mass spectrometry can be used to provide large numbers of measurements as both technologies continue to advance [13][14]. As unprecedented amounts of same-animal neural, metabolic and immune data come forth, it is necessary to not only have algorithms that make inferences from the data, but to also have mechanisms to represent the data in a manner through which insightful questions can be asked. We aim to establish an analytical framework that links brain state transitions with heterogeneous data collected at different timescales from disparate systems. A point of this work is to motivate matrix-based methods that can provide a unified comparison between measures obtained at single cell resolution but with no electrophysiological correlate, and measures obtained from bulk activity in conjunction with EEG/EMG. A time-hierarchy-modulation (THM) matrix is introduced in light of powerful experimental tools that enable us to record neural, metabolic, and immune signals and correlate how their activity leads to behavioral states such as consciousness. It will be intriguing to explore the predictive capability of such techniques in determining phenotype or behavior based on cellular activity from the various systems in an organism.

## 2. Necessity of a Time-hierarchy-modulation (THM) Matrix

Conclusions may be drawn by examining spatio-temporal dynamics associated with cellular activity when considering data in matrix form in concert with real-time behavioral monitoring. Powerful experimental tools are allowing for additional temporal and spatial resolution by delineating populations of neurons that are believed to be involved in complex behaviors such as sleep and consciousness. The tools promote a presentation of increasingly more tangible and ambitious models for quantifying the response of metabolic and immune signals in addition to neural responses and how the activity levels communicate to result in a brain state. Acknowledging the complexities of consciousness, one limitation incorporated in this presentation is restricting the space of brain states to two values. As we apply the framework within the sleep regulatory system the two values will be equated to sleep and wake/arousal. It is well-known that neither sleep or consciousness can be effectively partitioned into a few modes from a neural, metabolic, or immune perspective. This is done for ease of presentation to introduce the desired notions and does not cause a loss in generality. For instance, an example of how the analysis may be extended would entail the space of activity that leads to sleep being bifurcated for REM and NREM. The incorporation of different gradations of the two most salient brain states is an important extension that will complicate the model, but this is understandable when considering data attained from freely-moving animals.

The understanding of alterations in neural activity between various portions of the brain is usually quantified by pairwise correlations. With progress in measurement techniques (e.g. multifiber photometry) and the ability to attain higher-dimensional data from multitudes of locations in animals, increasingly more sophisticated techniques for quantification are being sought [15]. Dimensionality reduction techniques such as principal component analysis (PCA) and its variants have found ubiquitous use on neuroscience datasets attained via continuous recordings from large numbers of neurons. Such data is presumed to have relatively low dimensionality either via post-hoc analysis or based on prior findings. However, when considering neural data that has been collected via methods that operate on different timescales in unison with metabolic and immune data, it is not immediately obvious whether the behavioral state can be identified via lower dimensional subspaces. The notion of a THM matrix is presented as an intra- and inter-system means of representing and interpreting collected data. A goal of the analysis is to conjoin data from distinct systems and at different recording times to attain a unified view of monitoring and predicting a brain state. The time, hierarchy, and modulation aspects of a THM matrix shall now be discussed to substantiate the idea. Consider multi-dimensional recordings made from an animal at each time instant – we use the symbol **X** to denote the THM matrix and assume a dimensionality of *T* × *M*. The number of rows T constitute the temporal samples collected while the number of columns M denotes the output of the recording units. After successive time instances **X** may have a realization given by

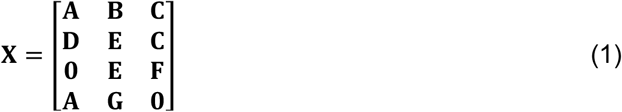

with the constituents **A, B, C, D, E, F, G**, and **0** denoting sub-matrices of smaller dimensionality. The **0** is used to denote a matrix or vector consisting of all zeros. The above realization of **X** indicates the consideration of three recording units across four successive time intervals. For example, in the case of the THM matrix containing neural data, the three column submatrices of **X** may correspond to activity from the tuberomammillary nucleus (TMN), locus coeruleus (LC), and lateral hypothalamus (LH) during a recording. It should be noted that the aforementioned component matrices that reflect the activity of various circuits may have vastly different dimensions. For instance, it is possible that **A**: 20 x 10 and **B**: 20 x 200. It is necessary to derive conclusions from a THM matrix either after or during the data collection phase. This will be done via a function f(.) that serves as a means of identifying differences in **X** matrices obtained from animals that are genetically or pharmacologically perturbed versus matrices constructed from healthy controls. The function f(.) can be concocted to provide the categorization **X** → *f***(X)** → brain state, and may be a classifier that has its parameters optimized via THM matrices that form a training set prior to f(.) making predictions from THM matrices that comprise a test set.

The presence of hierarchy in a THM matrix starts by considering its decomposition into constituent submatrices that can be used to assess the relative importance of groups of cells, genes, or other entities that are being recorded from in the induction of a behavioral state. Consider the system

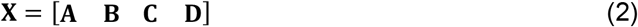

with *f*(**X**) = 1, if the activity in the component matrix **A** has been categorically manipulated to attain **A**=**0**, then the system reduces to the following THM matrix

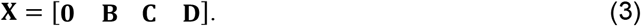

If *f*(**X**) = 0 and it can be further observed that 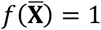 with 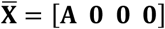, then **A** will be said to have hierarchy over **B, C**, and **D** in the sense of f(.) inducing brain state “1”. It is also possible to use the THM matrix to probe the redundancy of the circuitry in promoting a brain state. For instance, consider (2) and the problem of investigating whether there exists a 3-tuple of matrices **M**_l_, **M**_2_, **M**_3_ from the set {**A, B, C, D**} that lead to *f*(**A, B, C, D**) being adequately approximated by *f*(***M***_l_, ***M***_2_, ***M***_3_), e.g. if *f*(**A, B, C, D**) ≅ *f*(**A, B, C**) in which case the activity of the fourth recording unit may be deemed as redundant within the context of the brain state studied. From an experimental perspective, the activity of the fourth recording unit may have been manipulated via pharmacology or lesions. In fact, the incorporation of modulation in the THM matrix follows from considering how exogenous manipulations can affect the matrix structure and whether the resultant matrix leads to an alteration (i.e. modulation) in brain state. The silencing of the activity of **A** in (3) has been discussed via **A**=**0**. While this is an extreme occurrence, the modulation is a change in the activity of **X** and the fact that it may affect the output of f(.). The considerations of hierarchy and modulation become more involved in the realistic scenario of the THM matrix being time-varying. For instance, for the THM matrix shown in (1) the third recording unit takes three different values during the recording – it may be that a submatrix exhibiting hierarchy at an earlier epoch in the recording does not express the same dominance over the other submatrices at a later epoch. Examples of this are pervasive in accounting for circadian effects where neuronal and metabolic signals alter their degree of importance in the induction of a brain state at different time points in the cycle.

From a practical perspective, the matrix **X** must be constructed via an algorithm with inherent rules and restrictions. Even in the scenario of the behavioral or brain state presumed to take on a scalar value, it is unreasonable to concoct solving the optimization problem

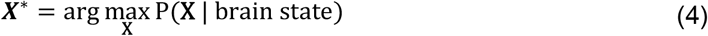

without the problem assumed to have structure such as its components being binary, sparse, or Gaussian. Furthermore, we would not anticipate the solution of an optimization problem such as (4) to be unique, and it would be difficult to vet among the solutions. We shall say more about (4) when discussing the notion of stability where an analytical reference point such as **X\*** is desired. Thus, **X** may be specified based on a heuristic that is amenable to the practicalities associated with the data collection and brain state monitoring system. A more practical, data-driven protocol for forming the THM matrix should encompass the following steps.

<u>Step 1</u>) Specify M, the number of recording units under consideration. The measurements in the units will comprise the columns of the THM.

<u>Step 2</u>) Determine the contribution of each of the M components in a row of the THM matrix to inducing the brain state that is being scrutinized. This should involve the normalization of the components with respect to a reference value such as the maximum activity level possible.

<u>Step 3</u>) Specify the temporal resolution, or synonymously the timescale at which successive samples of data are recorded. The number of rows in the THM will be determined based on the number of temporal samples/measurements taken.

Figure 1 depicts an outline of the steps in constructing a THM matrix. Irrespective of the timescale considered in forming **X**, it is unrealistic to expect a memoryless or Markovian structure among the successive rows of **X**. Rather, it is more feasible to explore the number of prior rows, i.e. **X**(i - 1,.), **X**(i - 2,.), …, **X**(i - L,.) that affect the formation of row **X**(i,.). The investigation of such question probes the degree of memory, L, within the system. This notion can be more formalized via the relation

**Figure 1:**
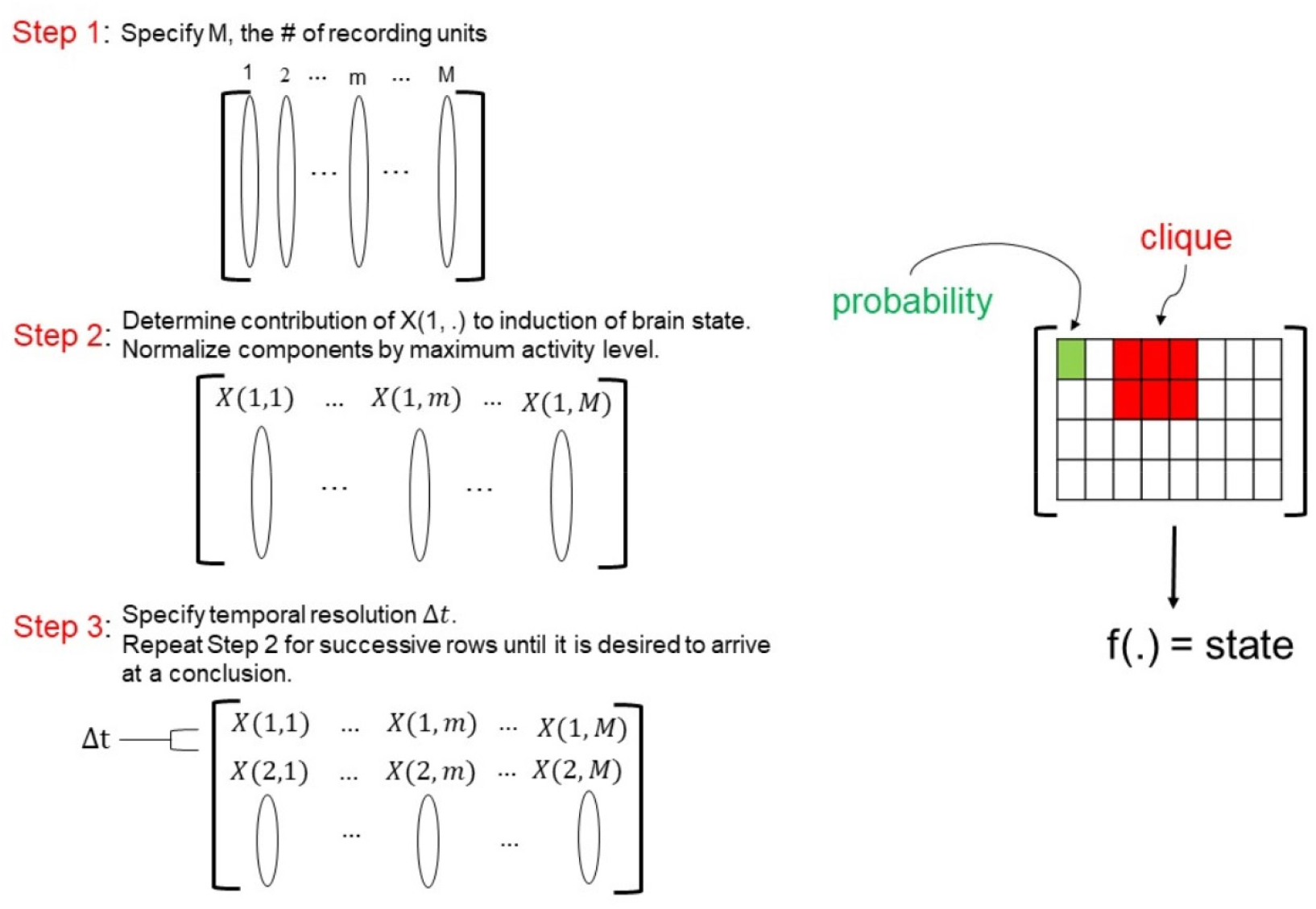
An overview of the time-hierarchy-modulation (THM) matrix notions within the context of studying how brain, metabolic, and immune system activity is modulated by consciousness. Left: The general steps involved in forming a THM matrix. Right: The units comprising a THM matrix. Each matrix component is a probability value that has been attained via a normalization. The sign of the component denotes whether the activity of that component promotes (+) or impedes (-) arousal. A clique represents successive columns of a THM matrix that are similar enough in activity, anatomy, or other specified properties to be considered as a collective entity. The function f(.) denotes a mapping from a THM matrix and a qualitative value (i.e., wake or sleep). Such a function may be a classification rule that has been formulated from prior realizations of the THM matrix.

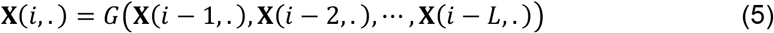

and perhaps approximated in the simplified scenario where G(.) is assumed to be linear via

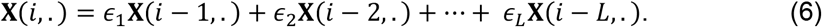

The weights {*ϵ*_l_, *ϵ*_2_, ⋯, *ϵ*_L_} may be learned and updated via a regression or artificial neural network technique as it would not be necessary to include any regularization constraint. We discuss tracking and prediction of brain state via a THM matrix. Following successive recordings, S consecutive rows of a THM matrix are defined as a “block” that is deemed temporally long enough to reflect the integration of the measurements. A relatively simple instance of the THM matrix being used to predict a brain state entails the evaluation of the following conditions at times i = 0, 1, …, T-1. For the ith block consider a threshold event #1 (TE-1) defined by

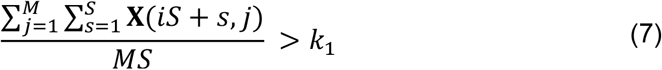

and the threshold event #2 (TE-2) defined as

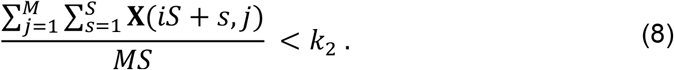

It is valuable to equate the events TE-1 and TE-2 to be prospective brain states. Since the two events are dependent on the thresholds k_1_ and k_2_, techniques such as point estimation [16] should be used to determine these two values that correspond to transitions in brain state (e.g., sleep to arousal). In fact, the structure of a THM matrix will enable the matrix to be partitioned with components being studied and relationships investigated among the probed recording units. A choice for the function f(.) has been addressed in (7) and (8) via

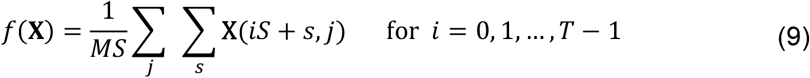

in conjunction with T blocks and the thresholds k_1_ and k_2_. The above choice of f(.) is certainly not a unique specification since one may select other collective operations on the elements of **X** to correlate with brain state. Akin to a block, we will use a “clique” to denote columns of a THM matrix that are similar enough in activity, anatomy, or other specified properties to be considered in unison. The selection of a clique is to some degree arbitrary, but it should follow from an anatomical facet or experimental constraint. We believe that the consideration of cliques will be valuable in portraying interactions that take place among the neural, metabolic, and immune systems. It will also allow existing graph-theoretic and newer edge-centric [17] views of network analysis to be applied on datasets in ways that have not been conceived in the past.

## 3. THM Matrices for the Sleep-regulatory System

Since the presented analysis is geared towards the sleep-regulatory system we shall consider the components of a THM matrix to reflect the likelihood of that component inducing arousal. More specifically, the elements are taken to be probabilities via the assignment **X**(*i, j*) ∈ [− 1, 1] with the sign associated to each component designating whether the magnitude reflects the degree to which the unit promotes (+) or impedes (−) arousal. We describe the formation of separate THM matrices for EEG, calcium imaging, metabolic, and immune data collected from mice within the behavioral context of sleep and arousal. The output of each of the four THM matrices may be determined by four different f(.) functions with the output of a function denoting a “state” for the THM matrix. Figure 1 depicts the relationships that we have discussed among the components of a THM matrix. There are similarities in the construction of the THM matrices, thus increased discussion will be given to the first matrix which will reflect EEG activity.

A spectrogram contains the quantitative information necessary to form a THM matrix for EEG recordings. Define a THM matrix for EEG data as THM_EEG_ with dimensionality *T*×*F* where F denotes the number of frequency bins with available power levels, and T is the temporal window of recording. We shall consider the matrix

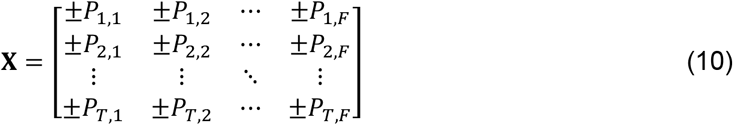

with the elements being probabilities assigned to the power level of the frequency bin and the component-wise +/− sign dictating whether the power level at that time-frequency promotes/impedes arousal. For instance, P_i, j_ = 0.7 may correspond to a recording from a frequency bin in the gamma band (30-80 Hz) with the positive value representing a high power level having a likelihood of inducing arousal. In order to have the components of THM_EEG_ be probabilities the matrix values would be normalized by the maximum power level attained during a recording. The effects of the sleep/wake cycle on the probabilities are captured in the rows of THM_EEG_ as we expect P_i, j_ to increase the longer during a recording that an animal has remained awake. Following the population of THM_EEG_, a sensible construction would encompass the various frequency bands such as theta, beta, and gamma that comprise adjacent columns each being a clique. Works such as [4] have discussed the non-Markovian nature among what would constitute the rows of THM_EEG_. With the relative availability of EEG data across various animal models, it may not be difficult to compute estimates for the weights in (6) via standard regression techniques.

From an algorithmic perspective, developments in calcium imaging methods for recording from neural populations present exciting avenues. Works such as [18] [19] [20] have discussed algorithms that draw upon the conflation of matrix analysis, statistics, and machine learning (ML) in order to identify neuronal properties such as the neurons’ precise coordinates. Matrix-based techniques may also be exploited to quantify the spatiotemporal traits of GCaMP and fiber photometry data that have been correlated to brain states. The dynamic contribution of calcium activity in different neural circuits or cell types to a brain state can be captured via the *T*×*M* matrix THM_Ca_ where M denotes the number of neural populations which may span several brain regions, and T is a temporal window of observation. The elements of THM_Ca_ represent probabilities associated with calcium levels at various neural populations and their sign indicates whether their activity promotes or impedes arousal. Following the construction of THM_ca_, a logical consideration would encompass each neural circuit being a clique.

Works such as [6] have provided models that are suggestive of the interplay of metabolic processes that are either upregulated or downregulated during sleep with the purpose of optimizing energy conservation. In fact, this is argued to be ubiquitous among species and across evolution. It is important to have measures that allow one to draw inferences between metabolic data and brain-state since the systems are interlinked. For instance, with respect to arousal, decreased levels of brain lactate have been implicated in numerous studies [21][22]. A THM matrix for the metabolic cell data is defined as THM_M_ with dimensionality *T*×*M* where T refers to the number of measurements (i.e. the time variable), and M denotes the number of recording units whether it be voxels or metabolic concentration levels. In fact, it is exciting that high-throughput PET/CT imaging architectures such as [23] are providing advancement to the acquirement of in vivo metabolic data by providing brain images over an hour recording interval. Additionally, works such as [24] have considered microdialysis and EEG recordings collected simultaneously from the medial prefrontal cortex (mPFC) and primary motor cortex (M1) in mice. A minimum of 240 microdialysis samples were collected from the two regions of mice in three cohorts. The microdialysis portion of such data can be represented, analyzed, and compared via 41 x 480 THM_M_ matrices since 41 time-points were collected from each animal in 15-minute increments. Whether monitored via voxels or metabolic concentration levels, we shall consider a clique in THM_M_ as a metabolite that has its activity level tracked. Lastly, it is possible to consider THM_M_ being formed from BOLD recordings with the ROIs corresponding to the columns of the matrix. Although the signal is collected from the brain, the metabolic measure of BOLD lends it appropriate for THM_M_.

A THM matrix for immune cell data is defined as THM_IC_ with dimensionality *T*×*I* where T denotes the number of recordings and I denotes the number of different immune markers with relative expression levels available. The immune and arousal systems have been discussed as having bidirectional interactions. For instance, we recently showed that optogenetic stimulation of corticotropin-releasing factor (CRF) neurons in the hypothalamus activated by stressful stimuli leads to a coordinated response in the periphery that includes both innate and adaptive immune compartments [25]. We attribute this as providing evidence for a clique in THM_EEG_ associated with recordings from the hypothalamus having interactions with entries of THM_IC_ associated with CyTOF recordings from MHCII+ dendritic cells of the innate immune system and B lymphocytes of the adaptive immune system. There is also a growing body of literature studying cytokine and interferon signaling on sleep modulation [26]. For instance, TNF and IL-1 have been discussed as keeping track of arousal history and acting as threshold-based contributors to the induction of sleep [27]. Furthermore, the role of sleep modulators such as hypocretin are being increasingly considered in the reciprocal interactions that exist among sleep, metabolism, and immune function. It has also been suggested that leptin inhibits the production of hypocretin, and that hypocretin antagonism improves sleep quality in association with improved glucose metabolism [28]. Such results are in agreement with the hypothesis that the dysfunction of hypocretin producing neurons alters glucose concentrations via the sympathetic nervous system [29]. In considering data provided from immune markers as the method for population THM_IC_, current CyTOF techniques allow for I=40 dimensions [30]. Following the construction of a THM_IC_ it would be sensible to equate each marker to a clique.

A high-level view of the formation of THM matrices and the determination of a brain state from each THM matrix is provided in Figure 2. The conditions in (7) and (8) may be evaluated in real-time during a recording session, as the specifications were made under the premonition that sustained bursts or silences in activity by the recording units that comprise THM_ca_ and THM_EEG_ are predictive of arousal. In the case of THM_I_ it may be that S=1 due to the larger time interval in-between measurements. In fact, it is necessary to differentiate among a behavioral outcome being threshold-based per time-instant (i.e. when S=1) or being determined after an interval (i.e. for S > 1). The two analyses are not at odds and can be concurrently pursued via the same THM matrix. It should be apparent that the THM matrices discussed above evolve on different timescales. EEG and calcium imaging data typically provide millisecond resolution whereas metabolic and immune recordings may be made on intervals that are interspersed by days. The THM matrix analysis is designed to address this issue since each matrix is formed on its own timescale with conclusions derived from THM matrices not being directly dependent on one-another. The interaction among the cliques of the four THM matrices will need to account for the timescales among the matrices. For instance, with respect to the non-Markovian relation suggested by (5), it is expected that L would be smaller when considering the rows in THM_IC_ as opposed to rows in either THM_Ca_ or THM_EEG_ that have been collected under similar experimental conditions. This is because the temporal dependence among successive EEG or Ca-imaging recordings are expected to be greater than those among consecutive recordings from immune cells since the latter would be further separated in time. As previously mentioned, the reflection of a THM matrix on brain state is accounted for via a state or outcome derived from the entire matrix. A unification of the outcomes attained from the THM matrices to make a decision on the brain state will be discussed in the ensuring sections.

**Figure 2:**
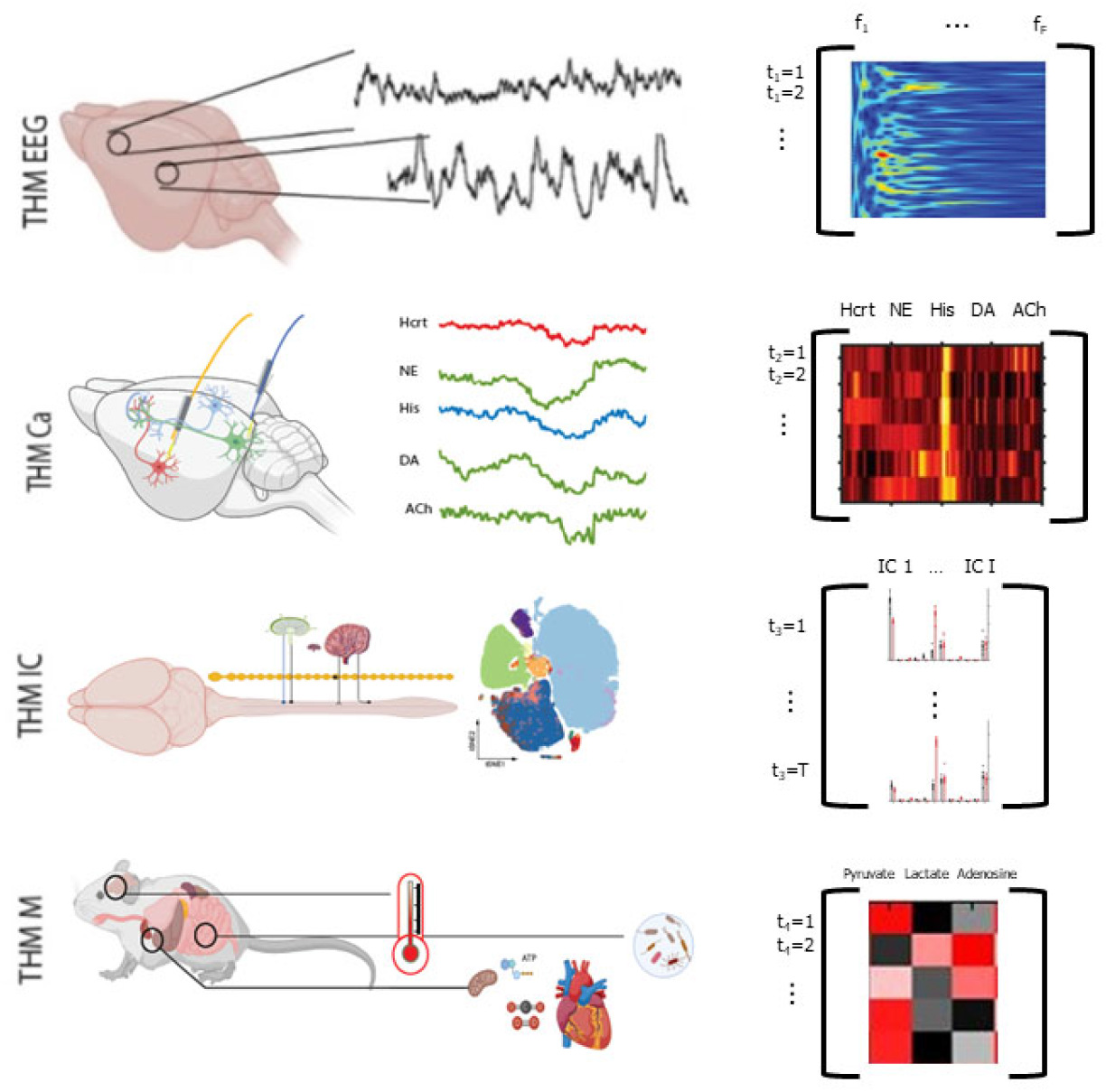
A consideration of recordings from the central nervous, metabolic, and immune system leading to the formation of respective THM matrices. The rows for each of the four matrices correspond to successive measurements while each column represents a recording unit.

## 4. States and Stability Analysis in the Sleep-arousal System

The elements comprising a THM matrix have been discussed as being probabilities. A clique or a group of components that interact with other cliques in the same matrix or with cliques in other THM matrices are meant to reflect the presence of pathways or subnetworks. The interaction of states and dynamic connections within a complex system that may be prone to dysfunction leads us to consider fundamental analytical notions. Stability is a central theme in dynamic system theory and encompasses long time horizons and principles within a constructed system where gains can be adjusted, exogenous signals are applied, and feedback from various points may be reintroduced into the system. The stability of the system is then evaluated via the properties of a matrix characterizing or constituting the system. Although empirically-motivated, and containing information directly from the probed systems, the THM matrices that we have discussed do not possess structure in the matrix-theoretic sense (e.g. symmetric or Toeplitz). We reformulate stability when considering a network designed by nature where engineering is not readily achievable, and consider the system as being stable or unstable with respect to a reference. The THM matrix analysis will consider a continued deviation of collected matrices from the reference as being an instability. Such a notion is not asymptotic in the system-theoretic sense of stability analysis, but neither are the timescales considered in the nervous, metabolic, or immune systems during recordings.

Consider a THM matrix **X** and a corresponding reference THM matrix **R** that has been attained while the animal was in the same brain state under “healthy” conditions. A sustained deviation between the two matrices can be quantified, and the system denoted as unstable if the difference is sufficiently large. One suitable analysis would involve comparing a norm between pairs of matrices to thresholds via

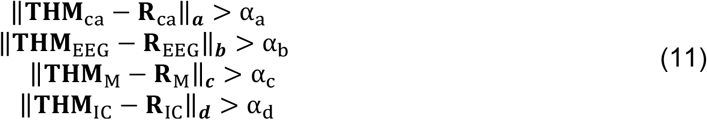

with α_a_, α_b_, α_c_, and α_d_ denoting the selected thresholds. A natural choice for the four norms in (11) would be the Frobenius norm. Nevertheless, one may choose to consider the 1-norm for the EEG data (i.e. b=1) because of its physical implication. Namely, the maximum absolute column sum of the deviation between **THM**_EEG_ and **R**_EEG_ would reflect that the expected power at a particular frequency has deviated from its reference threshold. The consideration of b=1 in this case instead of the Frobenius norm would follow from warranting the deviation of power at any one frequency band as being more important than the collective deviation in power across all recording frequencies in the EEG signal. Conversely, in the case of metabolic data it may be wise to consider the ∞-norm (i.e. c=∞) because the maximum absolute row sum of **THM**_M_ - **R**_M_ corresponds to a sudden change in metabolism at a particular time. The abrupt change can be due to immediate insulin production or consumption. The thresholds in (11) must obviously be adjusted in correspondence to the matrix norms that are considered. The fact that the thresholds will be determined via data leads us to believe that consensus may be reached on values after sufficient data collection across animals that are deemed similar enough with respect to genetic and phenotypic characteristics. In determining appropriate thresholds α_a_, α_b_, α_c_, α_d_ one possible heuristic would involve starting with somewhat high values such that none of the inequalities in (11) are satisfied. Through the consideration of successive four-tuples of measured THM matrices THM_EEG_, THM_Ca_, THM_M_, THM_I_ that have stemmed solely from aberrant or unhealthy animal recordings, the conditions in (11) should be satisfied at a high frequency (e.g. 95%). Accordingly, a threshold value in (11) may be iteratively lowered if its corresponding inequality is satisfied for a number of successive instances. The sequences of THM matrices from unhealthy recordings that are considered in the aforementioned procedure may be viewed as training data used for the tuning of the thresholds. Due to the inherent coupling among the four systems, several of the thresholds might be exceeded because of aberrant activity in one system. For instance, insulin resistance, diabetes, and obesity have been reviewed as being linked to and exacerbated by sleep disturbances or sustained arousal [31]. This is an example of a deviation in THM_M_ that will be affected by deviations in THM_ca_ and THM_EEG_. Thus, we consider an instability as being the more conservative occurrence of a simultaneous exceeding of all four thresholds.

The attainment of a reference **R** for each THM matrix has not been discussed yet. Determining such a reference matrix is not trivial and necessities us to revisit (4) with the aim of attaining a maximum a posteriori probability (MAP) estimate of the THM matrix for the considered brain state under non-aberrant conditions – i.e. equating **R** = **X**\* to be what is deemed acceptable for a healthy control. Assuming sufficient data has been collected from an animal during the brain state in question, one may conceive a discrete sample space to solve a more tractable version of (4) such as

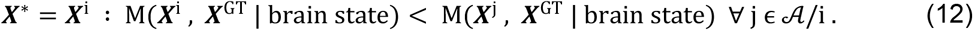

In the above ***X***^GT^ denotes a ground truth THM matrix that would be pre-specified and M(.,.) denotes a distance measure. In essence, (12) equates to evaluating pairs of possibilities, with respect to a measure M(.,.), in order to identify a THM matrix that is closest to ***X***^GT^. The solution **X**\* will as be used as the reference THM matrix **R**. The set *𝒜* in (12) contains the indices of THM matrices that are considered as candidates for being a reference **R** and may be expanded upon to contain more elements as more data becomes available. While the selection of a distance metric is not conceived to be difficult, the specification of a ground truth is a definite challenge that may be pursued through meta-analysis studies or via what is already known. In fact, ***X***^GT^ represents a generalized expectation for **R** in light of the animal breed, age group, weight, and other metrics that are not cohort specific. For instance, ***X***^GT^ may be standardized for large cohorts of genetically identical animals whereas **R** would be specified on a per cohort or even per animal basis. Figure 3 provides an example of the computations involved in selecting a reference from the ground truth and the measured THM matrices. In the case of THM_Ca_ with the use of a fiber photometry array that simultaneously records from brain regions consisting of DA, NE, Hcrt, and GABA producing neurons, we may attain the following THM_ca_ matrix after five time points

**Figure 3:**
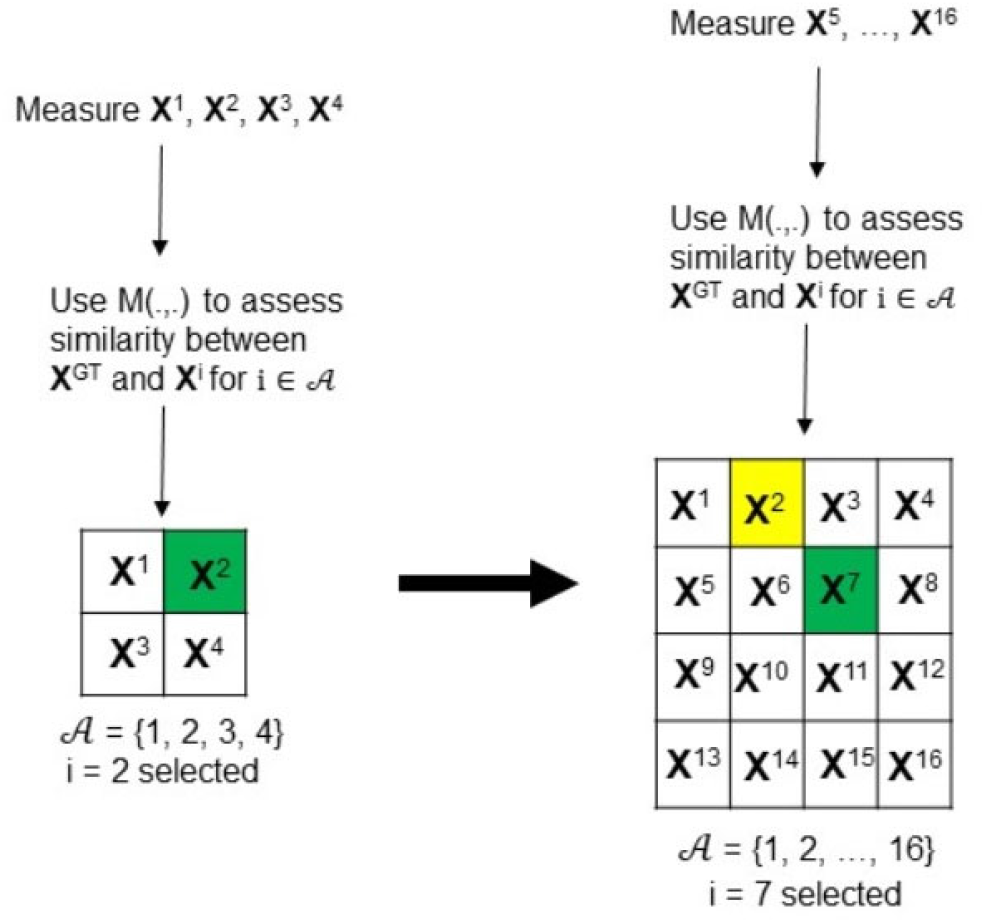
A view of how a reference THM matrix **R** can be iteratively specified for an animal or a cohort. Each **X**^i^ is a THM matrix that can be selected as the reference to which ensuing THM matrices of the same animal or cohort are to be compared in determining the stability of the system. As **X**^i^ : i = 1, 2, …, |*𝒜*| are collected for an animal under healthy conditions and in the brain state that is being scrutinized, evaluation of the discrete optimization problem in (12) will yield the THM matrix of that animal that is closest to a ground truth ***X***^GT^. A box colored green denotes the prospective THM matrix that has been selected as the reference following the comparison of the available data to a ground truth. This selection will serve as the reference **R**. In the example above, as twelve additional recordings of a THM matrix (i.e. **X**^5^, …, **X**^16^) become available, the prior reference – depicted in yellow – may be replaced by a new reference **R**=**X**^7^ that is even closer to the ground truth ***X***^GT^.

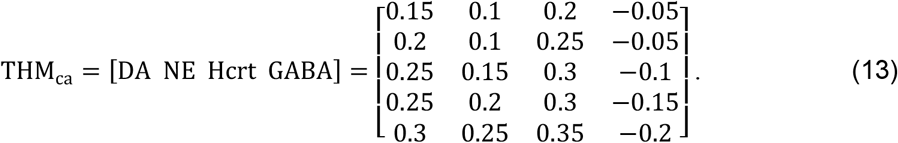

The above follows from the initial conjectures that the DA, NE, and Hcrt producing neurons are wake-promoting while the GABA producing population is sleep-promoting. The normalization by a maximum fluorescence signal level has also taken place in order to constrain the components to a magnitude that corresponds to a probability. In the scenario that the recordings are made at five time points per experiment, the successive measurements leading to 5 x 4 THM_ca_ matrices can be either used as prospective reference THM_ca_ matrices in the set *𝒜* or be averaged together to attain one prospective reference. The sequence of computing or using a previously-attained ***X***^GT^, and solving (12) to attain a reference **R** from a set of candidate THM matrices can be conducted, stored over the history of an animal, and be guided by a database where such information has been consolidated for animals with similar genetic and phenotypic traits. The evaluation of the stability analysis in (11) can be performed after estimated reference THM matrices and the thresholds are available.

As mentioned, the evaluation of the stability requires an ***X***^GT^ as a starting point. Fortunately, a ground truth for each of the four THM matrices can be hypothesized and then adjusted in iterative fashion following the application of unsupervised ML techniques that will look for similarity and trends in the data. There will also be a supervised ML component that will be implied in forming the ground truth – namely, the use of existing knowledge about the systems such as the posited roles of neural circuits, metabolites, and immune cells in inducing sleep or arousal. As the analysis is applied to increasing amounts of data and results are amassed from different labs, we expect that some degree of consensus will be reached on THM component values. It is beyond the scope of this paper to suggest more specific pipelines for forming each of the four THM matrices and provide data, but this is an avenue of current consideration.

## 5. Inter-THM Matrix Analysis via Network-Based Views

The consideration of EEG, multi-channel photometry, metabolic, and immune data derived from the same animal has not been conventionally undertaken due to challenges in its acquisition. It is known that the matrices correlate with arousal state as experiments continue to probe the interactions among the constituents in each realm. For instance, we have seen the increase in GCaMP photometry signals from Hcrt and DA producing neurons correlate with transitions from NREM sleep to wake [32][25]. While recording activity within various neural circuits is a central theme to neuroscience, it is being increasingly recognized that other systems should be incorporated in the discussion as they constantly adapt to environmental cues and to each other. As measurements of subcellular activity at different spatial and temporal scales are being made possible in the same organism, the analysis and models must consider that the byproducts of such activity is propagated along trajectories defined by both local and long-range connections. There is perhaps a necessity for models that account for the interactions between dynamics of different systems. Such modeling capability is crucial when attempting to understand a behavior as involved as arousal. A network view of the arousal system is a possible means to represent the degree of interaction among the cliques formed within a THM matrix and also among the cliques formed in the other three THM matrices. We shall consider the interaction of the matrices via a graph with nodes denoting cliques that have been identified in the neural, metabolic, and immune THM matrices, and the edges denoting the presence of adequate correlation among the respective nodes. Figure 4 depicts our presented view of inter- and intra-clique interactions among the components of the four THM matrices. The figure is meant to stress a network-based view of the coupling with the formation of cliques leading to subnetworks (or pathways) that exist within the system. Since correlations among neural populations can differ significantly with behavioral state, we expect network topologies to change for different brain states [33]. In fact, the edge coloring in Figure 4 is meant to distinguish among connections that exist during the two considered brain states. We have represented the system by an undirected graph to maintain generality – naturally, some interactions among cliques may be purely unidirectional. There are other caveats to the analysis as the absence of edges among cliques can not necessarily be equated to no strong interaction between cliques, but rather may be unknown or undiscovered. Nevertheless, we shall not assume or impose structure that might later be discarded as the methodology is applied to various datasets. Interestingly, in such a system the betweenness centrality of the cliques may be of more importance than other graph-theoretic measures as it would indicate a clique’s involvement in the most efficient – i.e. shortest length – paths. Of course, a measure of length would need to be formulated for the network based on multiple parameters that reflect the inter-workings of the neural, immune, and metabolic systems. The fitting of network structures to interacting cliques among the THM matrices will require data and perhaps iterations of model selection to arrive at possibilities. Such analysis is an exciting avenue that would warrant ensuing works.

**Figure 4:**
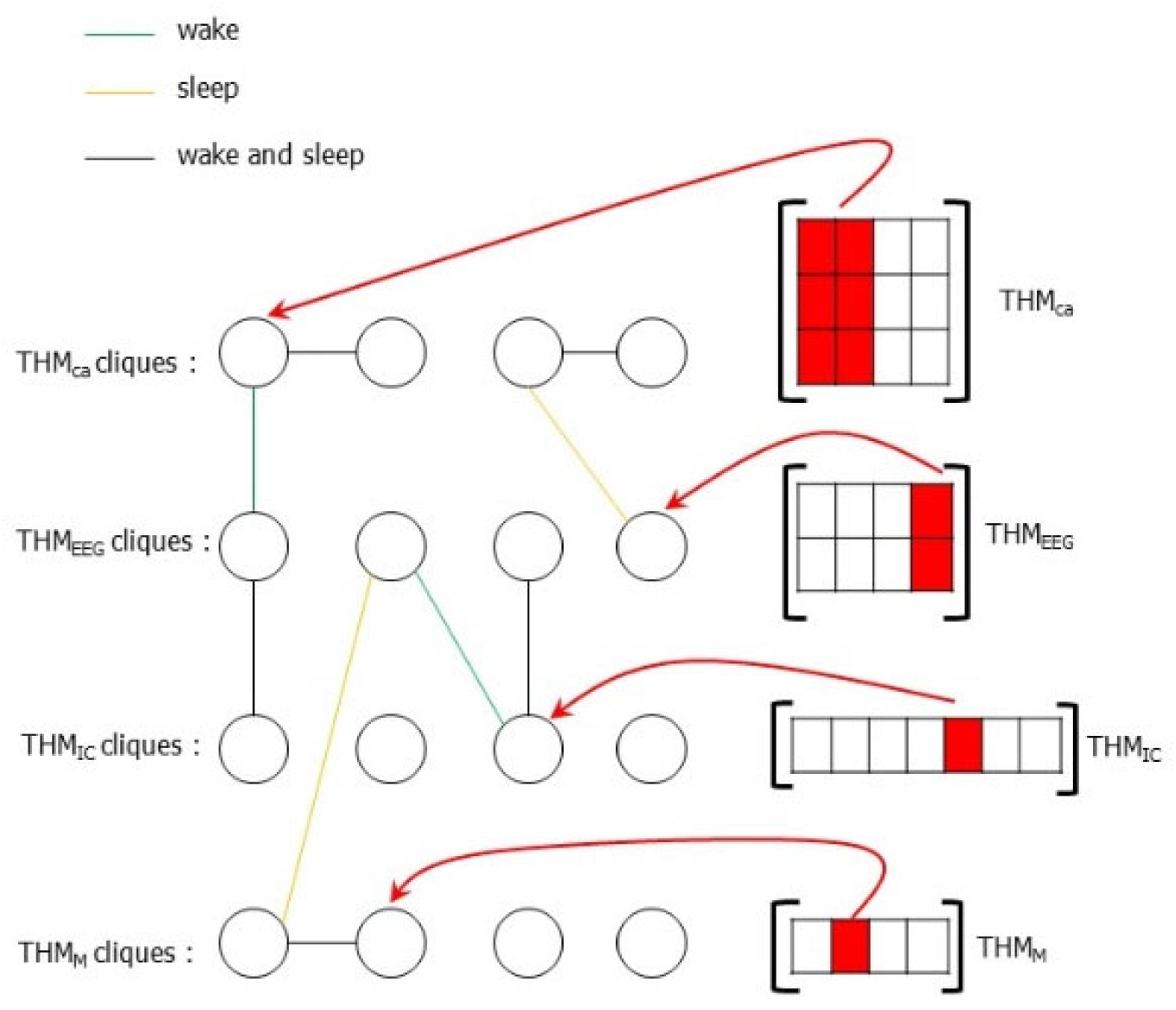
A network view of the interaction of states that are output from the THM matrices. The vertices denote the possible cliques that have been identified or specified from data collected for each THM matrix. The edge colors denote whether the clique interaction occurs during wake, sleep, or both states. The above depicts a time-invariant scenario where the cliques in the THM matrices are constant. This is a snapshot view whereas the more realistic time-variant consideration would encompass the network topology dynamically changing.

The brain has high metabolic needs, and for an arousal state, it would be insightful to consider THM_ca_ cliques as communicating with cliques in THM_M_ in order to form combinations that act in concert with, compensate for, or negatively correlate with one-another. As an instance of understanding how a brain state resonates to metabolic function, we have shown that a population of GABAergic leptin-responsive neurons provide inhibitory synaptic input onto Hcrt neurons in the hypothalamus [34]. Such a relationship would provide communication modeled via connected cliques in THM_M_ and THM_EEG_ as well as the same THM_M_ cliques communicating with cliques in THM_ca_. Compensatory responses between metabolic and neuronal networks have been discussed in works such as [35] where EEG recordings as well as metabolic measurements were taken in mice that experienced separate perturbations to their sleep pattern and diet. In fact, sleep disruption has been discussed as being both a driver and a by-product of disruptions to glucose metabolism [36][37]. Interaction of cliques in THM_IC_ and the other THM matrices also appear to be prevalent. An important mechanism of immune regulation by the brain includes the activation of noradrenergic neurons and norepinephrine release in the lymph nodes and spleen [38]. Furthermore, stimulation of the LH has been shown to result in an immunoenhancing response [39]. Interactions among the activity of Hcrt producing neurons in the LH and metabolites as well as immune cells were also studied more recently in [40]. The important question of whether the coordination of increased activity by DA-producing neurons along with greater expression of genes of interest in the immune system influence sleep coherence can be investigated by examining the interaction of the respective cliques in the THM_ca_ and THM_IC_ matrices.

An instance of the inter-network analysis is shown in Figure 5 as we consider findings from DA-producing neurons in the VTA. Metabolic data for this figure was collected in [41] while fiber photometry and EEG recordings were attained in [42]. The two connections from DA producing neurons of the VTA via THM_ca_ to THM_EEG_ cliques denote the inhibition of the power in the theta and delta frequency bands by the activation of the DA-producing neurons in the VTA during arousal. Since it is not currently known whether theta or delta EEG activity modulate the immune markers TLR4, CD25, CD80, and CD86; the corresponding connections between the THM_EEG_ and THM_IC_ cliques have not been included. It can be noted from the edge coloring in Figure 5 that we have only shown interactions that promote arousal. The discovery of clique dependencies and interactions that promote sleep are not as well understood and thus have not been shown. In utilizing collected data and the discussed framework, the choice of cliques may vary among different datasets, the interpretation of the data, and parameters used to fit the framework to the data. However, we expect the connections among cliques to be relatively consistent. The evaluation of altered connectively and communication as a result of the time-variant nature of the system is beyond the scope of the present work but an important extension.

**Figure 5:**
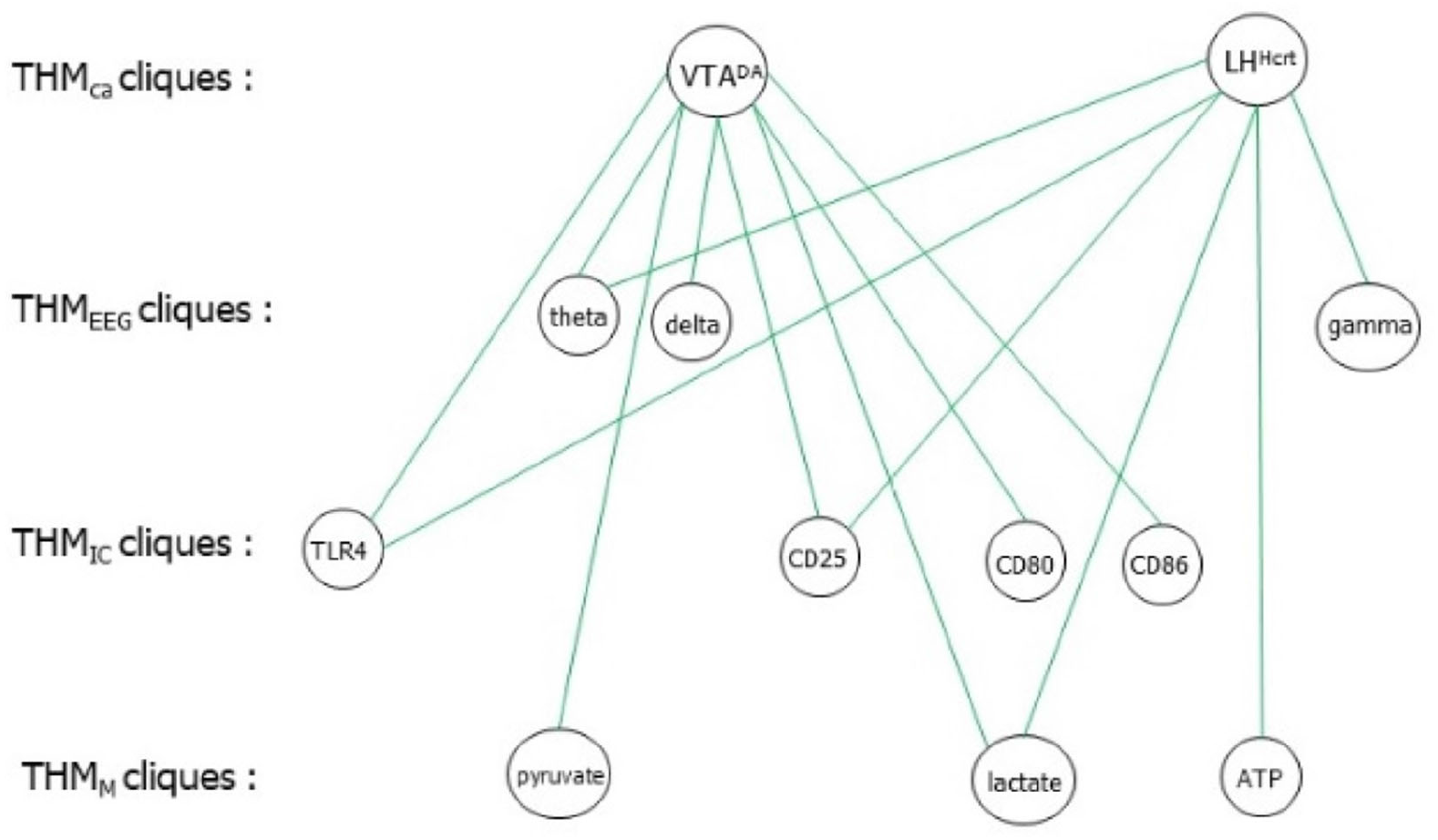
An example of inter-THM matrix analysis showing interaction among cliques. Although this is a snapshot view, it illustrates dependency among neural activity measured at different spectral bands via EEG, neural circuit-level calcium release via fiber photometry, markers associated with immune cell activity, and metabolic data. The clique interactions that involve EEG and fiber photometry data collected from DA producing neurons of the VTA were noted in [42] while the immune components were identified in [41]. The interactions shown among the metabolites, immune cells, and neural activity from the Hcrt producing neurons of the LH were determined in [40] while the interactions of the neural activity from fiber photometry with the metabolites were derived via in vivo microdialysis [24].

## 6. Conclusions and Future Considerations

The interaction of consciousness with the metabolic and immune systems throughout the sleep/wake cycle has been conceived for decades. However, the regulatory influence that each of these systems appear to place on one-another is only starting to be understood. The investigations have been aided by the development of instruments and experimental paradigms that enable heterogenous, multi-faceted data to be collected from the same organism over multiple time-points. This has constituted a relatively recent advancement in neuroscience since the tools that were used to provide a status of the nervous, metabolic, and immune system had been largely developed disjointly. From a data analysis perspective, protocols must be presented to convey and represent the data in a manner that is deductive. This has conventionally led to dimensionality reduction methods that rely upon sparsity existing in the volumes of data extracted from the different systems. Despite the successes of such techniques, there is a need to probe the data for structure that may be present in the underlying networks. This is especially true with the consideration of heterogenous datasets, i.e. neural, metabolic, and immune, stemming from complex networks where the inter-dependencies have not been completely discovered. Even exclusively in the realm of neural data collection, EEG and GCaMP present a challenge in constituting techniques that yield measurements at different timescales with each requiring a separate analysis for interpretation.

Matrix-based methods provide an efficient means for neuroscientists to compare the spatiotemporal neural activity to brain state. While it is important to consider experiments that provide data for the THM matrices, it is perhaps more crucial to identify the utility of prospective experiments within the scope of what has been mentioned. Advancement to already sophisticated technology is necessary to attain the data discussed herein. Nevertheless, progress is continuously being made on this front [43][44][45][1], and the incorporation of additional channels and optical fiber bundles to record from diverse populations of neurons will allow for further scrutiny and a more complete picture. For example, in the case of THM_ca_, it would be elucidating to study the simultaneous interaction of DA, Hcrt, NA, GABA, and 5HT producing neurons from the same animal with the cellular activity from the metabolic and immune systems as the animal transitions between wake, REM, and NREM sleep. Even within the subclass of GABA producing neurons, it would be valuable to attain simultaneous, separate-channel recordings from populations in the basal forebrain and the ventral medulla. With respect to interpreting data from the arousal system, a THM matrix will conjoin calcium imaging and EEG data collected from brain regions as well as metabolic and immune data to predict a brain state. The instantiation brings forth questions that must be addressed if experimental data is to be effectively incorporated into the framework. For instance, experiments that provide a consensus on the signs of the components in the THM matrices are necessary. Unfortunately, the majority of present experimental methods do not definitively distinguish among the portions of THM matrices that contain sleep-versus wake-promoting agents. Further targeted studies validating activity profiles that comprise each matrix will assist in ascertaining the sign and magnitude of the components. For THM_ca_, the DA and Hcrt releasing neurons that we have presented data for in the past are believed to be wake-promoting [42][32][25], and the recorded data from these respective populations would be accompanied with a positive sign in the THM matrix. In contrast, the activity from populations of GABAergic preoptic neurons that are associated with the onset of sleep would be assigned a negative sign.

The development of models that use optogenetic, pharmacological, or lesion analysis to delineate the sleep- or arousal-promoting properties of various populations of neurons was motivated in works such as [46] and remains an active area of research. In addition to calcium recordings of neuronal activity, a new wave of biosensors that allow millisecond scale determination of neurotransmitter dopamine [47] and transmitter/metabolite adenosine [48] are emerging and may improve the spatial and temporal resolution of the THM matrices. The possibility of such anatomically precise recording technologies for high-throughput data will permit various combinations of knock-out, pharmacological, or stimulation/inhibition scenarios to probe at the sleep/arousal function of each component in the system. It is important to have computational machinery in place to convey findings from the data as the experimental capability advances to provide increasingly sophisticated recordings. We anticipate the analysis to have multiple applications. While the identification of sleep disorders from the THM matrices based on the discussed notion of stability is a salient application, perhaps an equally interesting avenue is the discovery of brain states based on immune and metabolic responses. The interaction among cliques is a particularly important avenue to consider since it is being discovered that consciousness is a dynamic, multi-faceted phenomenon that encompasses several gradations. The ideas discussed in the present work have provided means to unite data from multiple systems. With respect to THM matrices, amassing and evaluation of data to find ground truths, evaluate the stability consideration and map interactions among cliques is formidable, but so is understanding consciousness.

